# Identifying single origin rare variants in population genomic data

**DOI:** 10.1101/2025.02.13.638116

**Authors:** Josh J. Reynolds, Vassiliki Koufopanou, Austin Burt

**Affiliations:** Department of Life Sciences, Imperial College London, Silwood Park, Ascot SL5 7PY, United Kingdom

## Abstract

Genomic analyses have shown that some mutations in large population genomic datasets may be the result of repeated, independent events at the same locus. However, the possibility of recurrent mutation is often ignored, even when it has the potential to introduce errors, such as when assuming co-ancestry for demographic analysis. Even rare variants such as doubletons, which should be particularly informative about recent demography, may have multiple origins despite arising relatively recently in the population. Here, we develop methods to (1) estimate the frequency of recurrent doubletons in a population genomic dataset from the occurrence of tri-allelic sites with two different singleton mutations, and (2) identify a subset of high confidence single origin doubletons based on the presence of a linked rare variant on the surrounding shared haplotype. Applying these methods to data for the malaria mosquito *Anopheles gambiae* sampled from across Africa, we estimate that ∼16% of doubletons had independent origins. We then identify a subset of doubletons highly likely (∼99%) to have a single origin, which consists of ∼68% of all the expected single origin doubletons (and ∼57% of all observed doubletons). The effectiveness of our methods is demonstrated by both further data analyses and coalescent simulations, and these doubletons are then used to test population genetic hypotheses about recombination, selection, and isolation by distance. The methods developed here should be useful for demographic inference when populations or sample sizes are large enough that recurrent mutation cannot be ignored.

## Introduction

A frequent assumption underpinning many population genetic models is that all copies of an allele in a sample descend from a single mutational event in a common ancestor in the genealogical past (Wakeley, 2008). Common frameworks that use this assumption include the “infinite alleles” and “infinite sites” models – that is, that each new mutation either introduces a new, unique mutant allele, or that each new mutation occurs at a new, unique position in the genome, and therefore there cannot be multiple copies of the same mutation arising from different sources (Kimura and Crow, 1964; Karlin and McGregor, 1967; Kimura, 1971). However, in reality some variants will be the result of the same mutation occurring on multiple branches of the genealogy, which are then inherited through successive generations into the present (Haldane, 1933; Jenkins and Song, 2011; Sargsyan, 2015; Wakeley et al., 2023).

The phenomenon of recurrent mutation has been most easily detected and studied at sites under strong, recent selection from new external factors leading to selective sweeps (Messer and Petrov, 2013; Smadi, 2017; Clarkson et al., 2021). Recurrent mutations at weakly selected or neutral sites have been less well studied, but if not accounted for may introduce errors into demographic analyses, with factors such as the incorrect inference of measures of relatedness between individuals and distortion of the site-frequency spectrum (Jenkins and Song, 2011; Seplyarskiy et al., 2021) leading to bias in population size estimates, measures of gene flow, and assessments of population structure (Borges, 2025; Schraiber et al., 2025).

Rare variants - those alleles found at very low frequencies in a population - are expected to be more recent and hence more likely to have a single origin than high frequency variants (Novembre and Slatkin, 2009; Mathieson and McVean, 2014; O’Connor et al., 2015). They may therefore be more geographically restricted, making them particularly useful for inferring recent, fine-scale demographic history, and distinguishing populations that are otherwise difficult to differentiate (e.g., O’Connor et al., 2015; Cubry et al., 2017). However, while the assumed status of a single origin may be appropriate for rare variants in some population genomic datasets, in others multiple origins of the same mutation may be more common than is widely assumed (Mathieson and Reich, 2017; Seplyarskiy et al., 2021). These may include highly diverse species that have large populations or those with high mutation rates which contribute to an excess of new mutation entering the population at polymorphic sites (Borges, 2025), and those which have undergone recent population expansions (Wakeley et al., 2023), especially if a great many samples have been taken - an increasing occurrence with modern sequencing approaches (Schraiber et al., 2025). Demographic inference in these species would be improved by having reliable methods that allow one to identify a subset of rare variants that are highly likely to have a single origin (Johnson et al., 2022).

In this paper we focus on the rarest of variants that can be useful for indicating relatedness, doubleton mutations, which occur in only two individuals in a population genomic dataset (Mathieson and McVean, 2014). We first describe an approach for estimating the proportion of doubletons that have multiple origins, based on an analysis of triallelic sites. We then outline how one can update the probability that a specific doubleton has a single origin based on the presence of another linked rare variant. We apply these methods to both coalescent simulations and published data from the *Anopheles gambiae* species complex (henceforth *An. gambiae* sensu lato, or s.l.), the primary vector of malaria in sub-Saharan Africa and therefore responsible for hundreds of thousands of deaths every year (The Anopheles gambiae 1000 Genomes Consortium, 2017, 2020; World Health Organisation, 2020). Finally, we use single origin doubletons from *An. gambiae s.l* to test population genomic hypotheses involving recombination, selection, and isolation by distance.

## Theory

For doubletons in a large dataset we will assume that the common allele is ancestral and either (1) the doubleton allele has a common origin and the two sequences are, at that position, each other’s closest genealogical relatives - i.e., they are reciprocal closest relatives (henceforth *RCR*); or (2) the doubleton allele arose twice at that site, independently, and the two sequences are no more closely related at that site than random pairs of sequences in the dataset (Fig. 1A). In principle there are additional possibilities, such as (3) that the doubleton allele is ancestral, or (4) that it is derived, with a single origin, but then there was a back-mutation to the ancestral state, leaving two sequences with the derived state that are not *RCR*, but these scenarios are expected to be significantly less likely than the other two and we will not consider them further.

**Figure 1:**
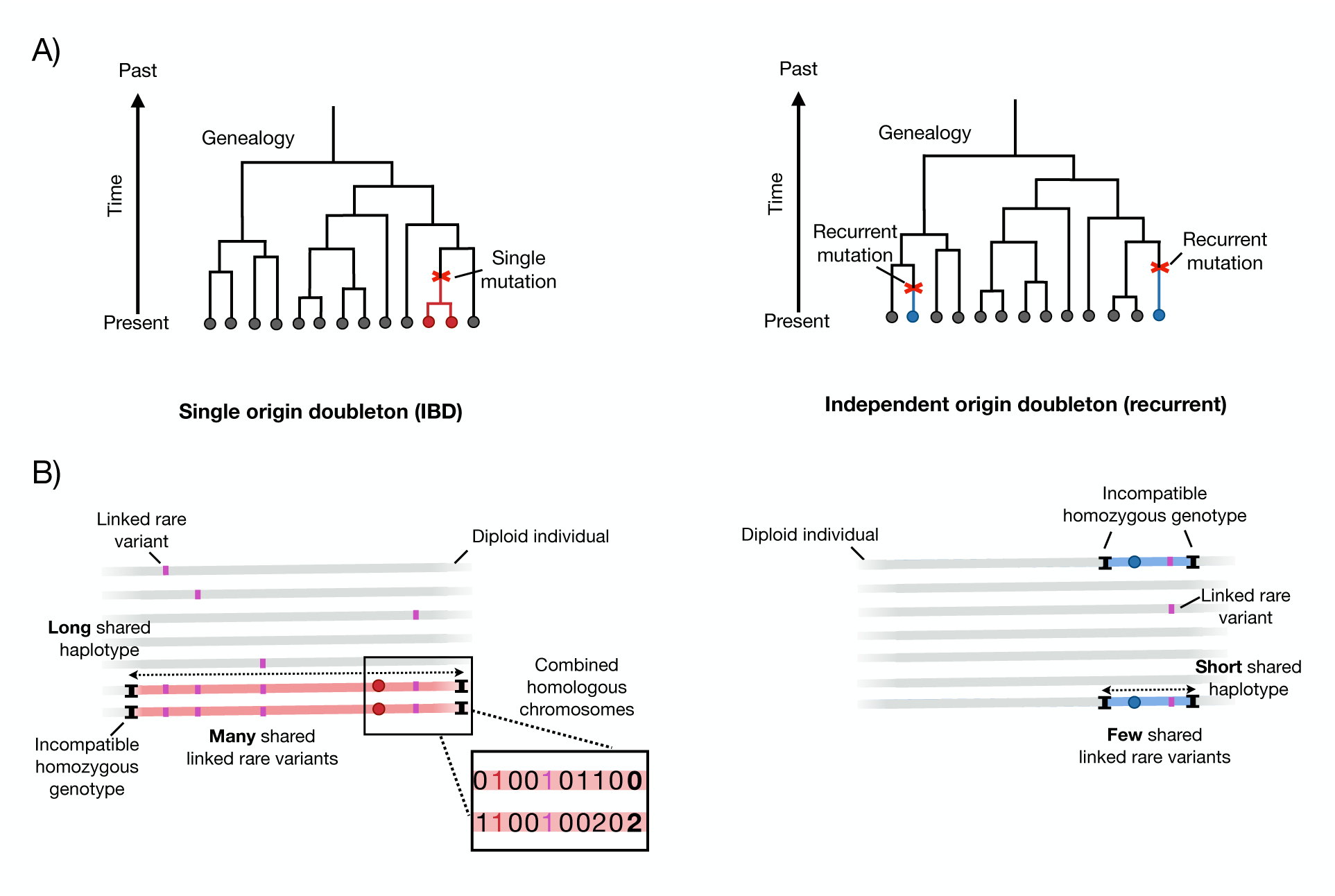
(A) The two most likely scenarios for a doubleton, either they have a single origin and the haplotypes are reciprocal closest relatives (*RCR*) at that site (left), or they have independent origins and the haplotypes are no more closely related than random (right). (B) Discriminating features of shared haplotypes around single origin and independent origin doubleton mutations. Example alignments of combined diploid homologous chromosomes. Haplotype boundaries are determined by identification of inconsistent homozygote genotypes between the two doubleton carrying individuals, either side of the focal mutation. Single origin haplotypes are generally longer, and share a greater number of other linked rare variants (purple) relative to those with independent origins. Inset in shows combined genotype data structure.

### Estimating the proportion of single origin doubletons

Our first goal is to estimate the prior probability that doubletons fit the first scenario above, that they have a single origin and the two sequences sharing the doubleton are *RCR* at that site, i.e., the conditional probability that the sequences are *RCR* given they share a doubleton, *P* [*RCR*|*d*]. To do so we use the rate of recurrent mutation at a single site, calculated from data on triallelic sites in which one base is found in all but two sequences, and then there are two different singletons (henceforth triallelic singleton sites). While, again, there are multiple possible scenarios that can lead to a polymorphism of this type, the most likely is that the common base is ancestral and there were mutations on two external branches of the genealogy to two different bases, with no mutations on any internal branch. Because the mutation rate from one nucleotide to another depends on the nucleotides involved (e.g., transition rates differ from transversion rates), we will estimate *P* [*RCR*|*d*] separately for each of the 12 types of mutation. However, for simplicity we will assume that the probability of a mutation of a specified type is constant throughout the genome - that is, that there are no neighbourhood effects of local sequence context on mutation rates.

Proceeding by example, suppose the ancestral state at a particular position is A. Let *P* [*A* → *C*], *P* [*A* → *G*], and *P* [*A* → *T*] be the probabilities that a mutation of an A on an external branch of the genealogy results in a C, G, or T, respectively, where the three probabilities sum to 1. Conditional on a site being ancestrally A and that there have been two mutations on external branches, the probability that one of the mutations is to a C and the other to a G is then *P* [*A* → *C, G*] = 2*P* [*A* → *C*] × *P* [*A* → *G*], and similarly for the other combinations, while the probability that both mutations were to a C (i.e., it is a recurrent doubleton) is *P* [*A* → *C, C*] = *P* [*A* → *C*]^2^. This last probability can therefore be expressed as

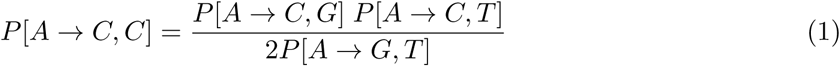

(see SI for derivation). These probabilities can be estimated from the observed occurrences of triallelic singleton sites in the genome, and the estimated number of recurrent *A* → *C* doubleton sites will be:

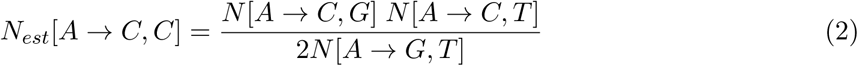

where *N* [*X* → *Y, Z*] is the number of sites in the genome for which the major allele is X with single copies of mutations Y and Z. Finally, if there are *D*[*A* → *C*] doubleton sites in which the major allele is *A* and the minor is *C*, then the estimated proportion that are single origin and *RCR* is

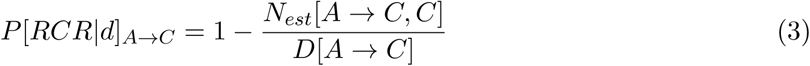

Analogous calculations apply for the other 11 types of mutation, and can be used to estimate the prior probability for each one that a doubleton has a single origin and is *RCR*.

### Estimating the probability of having a single origin

Our second goal is to identify a subset of doubletons that are highly likely to have a single origin and are *RCR*. We do this by updating the prior probability that a doubleton has a single origin using the presence or absence of a linked rare variant (*LRV*) near the doubleton. The use of *LRVs* is based on the logic that if the sequences are *RCR* at the doubleton site, then they are likely to be closely related at linked sites, and therefore have an increased probability of sharing a rare variant (Fig. 1B). Therefore, the presence of a rare variant should help inform on whether the sequences are *RCR*. The odds that a doubleton with a linked rare variant is *RCR* (defined as the probability it is *RCR*, *P* [*RCR*|*d, LRV*], divided by the probability it is not, 1 − *P* [*RCR*|*d, LRV*]) is equal to the prior odds that a doubleton is *RCR* (in the absence of information about linked rare variants) multiplied by the Bayes Factor (*BF*), which quantifies how much the odds of being *RCR* change due to the presence of a linked rare variant:

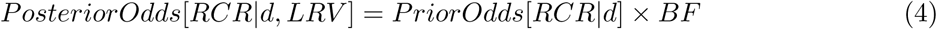

In the Supplementary Information we show that the Bayes Factor can be estimated by calculating the probability that individuals sharing a doubleton have a linked rare variant (*P* [*LRV* |*d*]); the probability that random pairs of individuals, which do not share a doubleton at that site, never-theless have a linked rare variant (*P* [*LRV* |¬*d*]); and the prior probabilities that the doubletons and random pairs are *RCR* (*P* [*RCR*|*d*] and *P* [*RCR*|¬*d*], respectively):

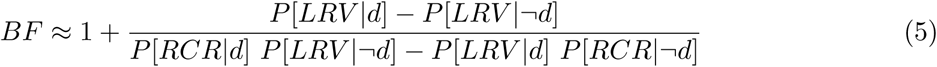

Thus we can calculate our probability of interest, that a doubleton with a linked rare variant is *RCR*, *P* [*RCR*|*d, LRV*], from the prior probability it is *RCR*, *P* [*RCR*|*d*], estimated as above, and from 3 other probabilities, which can be estimated as follows.

### Estimating *P* [*RCR*|¬*d*]

In a sample of *m* sequences, the probability that 2 of them chosen at random are *RCR* at a particular site in the absence of a doubleton, *P* [*RCR*|¬*d*], can be derived by considering the coalescence of lineages back in time. For them to be *RCR*, they must coalesce with each other before either one does with any other lineage.

Let 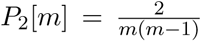 be the probability that the first coalescence event involves the two sequences, and 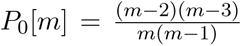 be the probability that it involves neither of them. If neither sequence is involved in the first coalescence, then the same logic applies to the second coalescent event, except there are now *m* − 1 lineages instead of *m*. The probability that *t* coalescence events occur before at least one of the two sequences is involved, and that the coalescence involves both sequences (and therefore they are *RCR*) is

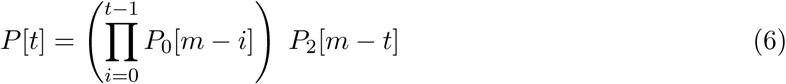

which can be summed over all possible values of *t* to get the overall probability they are *RCR*,

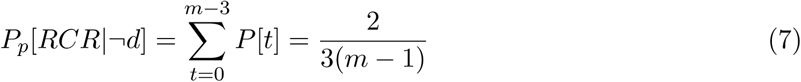

where the subscript *p* emphasises that this expression is for phased (or haploid) data. To avoid the uncertainties associated with statistical phasing of rare variants (Beck et al., 2025) we analyse unphased diploid genotypes in this paper. For two diploid individuals that do not share a doubleton there are 4 possible combinations of bases that could be *RCR*, and therefore to get the probability that at a particular site one of the sequences in one individual is *RCR* with one of the sequences in the other individual we must multiply the above expression by 4:

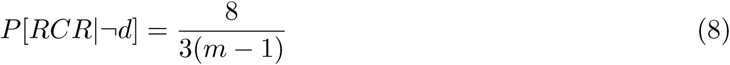

(In principle, for diploid individuals that share a doubleton, it is possible that the doubleton is not *RCR* but one of the other three pairs of bases between individuals is, but this alternative scenario will typically be very rare (i.e., *P* [*RCR*|*d*] *>> P* [*RCR*|¬*d*]) and we will not consider it further.)

### Estimating *P* [*LRV* |*d*] and *P* [*LRV* |¬*d*]

The remaining two terms in Eq. (5) are the probabilities of doubletons and random sites having a linked rare variant, *P* [*LRV* |*d*] and *P* [*LRV* |¬*d*], and we must now make precise the notion of having a ‘linked rare variant’. This could be done in multiple ways. Here we use a method that is appropriate for analysing unphased sequence data. In particular, we use the approach described by Mathieson and McVean (2014) to define a segment that extends in both directions from a focal base (doubleton or otherwise) until a site is found at which the 2 individuals are homozygous for different alleles (Fig. 1B). Following Mathieson and McVean (2014), we call these ‘inconsistent sites.’ These segments will typically provide an overestimate of the region shared by individuals over which no recombination has occurred. We then ask whether the segment contains (besides the focal base, which may be a doubleton), a rare variant that is found only in *n* sequences in the dataset. The probabilities *P* [*LRV* |*d*] and *P* [*LRV* |¬*d*] can be estimated simply by counting the proportion of doubletons and random sites that have a linked rare variant defined in this way. We repeat the analysis and calculation of Eq. (4) with different values of *n*, in order to achieve the desired level of confidence in a designation that a doubleton is *RCR* (e.g., 99% confident).

Note that sites that are or are not *RCR* will differ in the probability of having a linked rare variant both because the segments defined in this way are expected to be longer in the former than the latter and because the density of linked rare variants (i.e., on a per-base-pair basis) will be higher. Thus there are 2 contributing factors, which should help the discrimination.

## Results

### Identifying *RCR* doubletons

#### Dataset Description

We illustrate our approach using genome sequences from Phase 2 of the *Anopheles gambiae* 1000 Genomes Project (Ag1000G) (The Anopheles gambiae 1000 Genomes Consortium, 2020). We analysed variant data on the 42Mb chromosome arm 3L for 1,142 individuals of the sibling species *An. gambiae* sensu stricto (henceforth *An. gambiae* s.s.) and *An. coluzzii* collected from 13 countries across sub-Saharan Africa. Unphased data were used due to concerns regarding the ability of statistical phasing methods to accurately phase rare variants (Andrés et al., 2007; Snyder et al., 2015; Choi et al., 2018; Beck et al., 2025), and 3L was used as there are no reports of common segregating inversions on this chromosome arm (Coluzzi et al., 1985). The data were filtered to remove low quality sites or where any individual had a missing genotype, leaving 10,219,040 sites (see Methods for further details). This dataset contained 1,116,970 doubletons, and for each one we scanned the genome in each direction until finding a site at which the two individuals were homozygous for different bases, resulting in 754,006 unique haplotypes (i.e., some segments carried multiple doubletons) with a median length of 5.5 ± 0.009kb (SE). To calculate *P* [*RCR*|¬*d*] and *P* [*LRV* |¬*d*], a dataset of random or “non-doubleton” (¬*d*) haplotypes was created: for every doubleton we randomly chose two other individuals and, starting from the same site as the doubleton, again scanned out in each direction until finding a site at which the two individuals were homozygous for different bases, resulting in 1,115,755 unique segments. As expected, these randomly chosen haplotypes were considerably shorter, about half the length of those flanking doubletons (median 2.9 ± 0.003 kb).

#### Estimating the proportion of sites that are *RCR* (*P* [*RCR*|*d*] and *P* [*RCR*|¬*d*])

A total of 243,668 triallelic singleton sites (where one base is found in all but two sequences and then there are two different singletons at that site) were identified on chromosome 3L (Table 1A). Data from these sites were used along with Eqs. (2 & 3) to estimate the proportion of observed doubletons that are single origin and therefore identify sequences that are *RCR* (Table 1B) (for an example, see Supplementary Information). There is a 5-fold range in the estimated proportion of recurrent doubletons of different types, with C→T and G→A mutations having the highest proportion (∼25%), and A→C and T→G mutations the lowest (∼5%). Overall the estimated proportion of recurrent doubletons is 0.16, thus leaving *P* [*RCR*|*d*] = 0.84 doubletons as having a single origin.

**Table 1:**
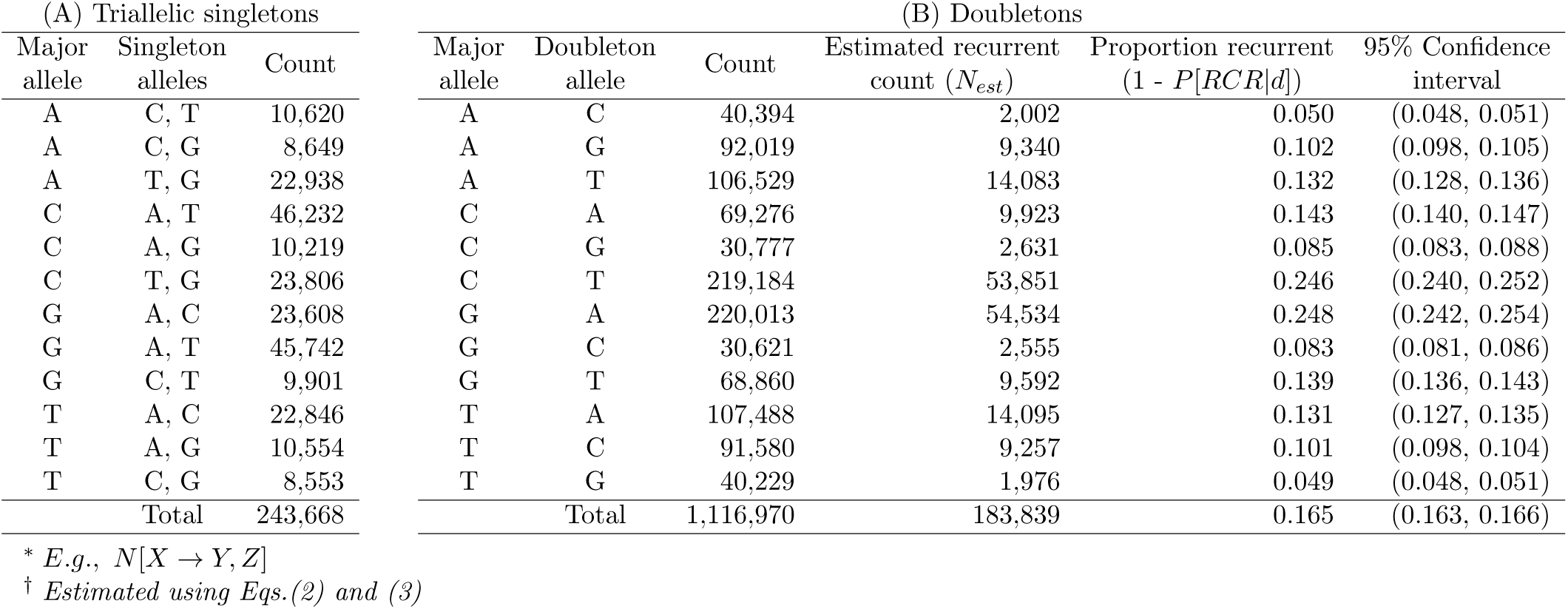
(A) Number of triallelic singleton sites^∗^, and (B) Number of observed doubletons and estimated recurrent doubletons^†^, on chromosome arm 3L.

| (A) Triallelic singletons |  |  | (B) Doubletons |  |  |  |  |  |
| --- | --- | --- | --- | --- | --- | --- | --- | --- |
| Major allele | Singleton alleles | Count | Major allele | Doubleton allele | Count | Estimated recurrent count ($N_{est}$) | Proportion recurrent ($1 - P[RCR\vert d]$) | 95% Confidence interval |
| A | C, T | 10,620 | A | C | 40,394 | 2,002 | 0.050 | (0.048, 0.051) |
| A | C, G | 8,649 | A | G | 92,019 | 9,340 | 0.102 | (0.098, 0.105) |
| A | T, G | 22,938 | A | T | 106,529 | 14,083 | 0.132 | (0.128, 0.136) |
| C | A, T | 46,232 | C | A | 69,276 | 9,923 | 0.143 | (0.140, 0.147) |
| C | A, G | 10,219 | C | G | 30,777 | 2,631 | 0.085 | (0.083, 0.088) |
| C | T, G | 23,806 | C | T | 219,184 | 53,851 | 0.246 | (0.240, 0.252) |
| G | A, C | 23,608 | G | A | 220,013 | 54,534 | 0.248 | (0.242, 0.254) |
| G | A, T | 45,742 | G | C | 30,621 | 2,555 | 0.083 | (0.081, 0.086) |
| G | C, T | 9,901 | G | T | 68,860 | 9,592 | 0.139 | (0.136, 0.143) |
| T | A, C | 22,846 | T | A | 107,488 | 14,095 | 0.131 | (0.127, 0.135) |
| T | A, G | 10,554 | T | C | 91,580 | 9,257 | 0.101 | (0.098, 0.104) |
| T | C, G | 8,553 | T | G | 40,229 | 1,976 | 0.049 | (0.048, 0.051) |
| Total |  | 243,668 | Total |  | 1,116,970 | 183,839 | 0.165 | (0.163, 0.166) |

\* *E.g.*, $N[X \rightarrow Y, Z]$
† *Estimated using Eqs.(2) and (3)*

To estimate the probability that our random pairs of individuals are *RCR* at a specified site, we use Eq. (8). With 2,284 haploid genomes in our sample, *P* [*RCR*|¬*d*] = 0.0012. Thus individuals sharing a doubleton are about 700 times more likely to be *RCR* at that site than a random pair of individuals.

#### Estimating the probability of sites sharing a linked rare variant (*P* [*LRV* |*d*] and *P* [*LRV* |¬*d*])

The probability a doubleton has a linked rare variant, *P* [*LRV* |*d*], is given by the proportion of doubletons with at least one rare variant in the flanking segments out to the nearest inconsistent homozygote sites. Figure 2A shows the fraction of doubletons with an *LRV* on their flanks as a function of the frequency of the *LRV* in the dataset (shown both as cumulative counts, where doubletons are included if they have a linked variant present in anywhere from 2 to *n* copies in the dataset, and as incremental counts). As can be seen, the incremental values trend downwards, with proportions of higher-count *LRVs* becoming increasingly smaller while the cumulative counts increase at a diminishing rate. This same analysis on the random pair haplotypes shows a relatively flat incremental plot and the cumulative plot is therefore linear (Fig. 2A). Interestingly, random sites are also less likely than doubletons to have a linked singleton site (32% vs. 66%), presumably because the segments out to the nearest inconsistent homozygote site are longer for doubletons than for random pairs.

**Figure 2:**
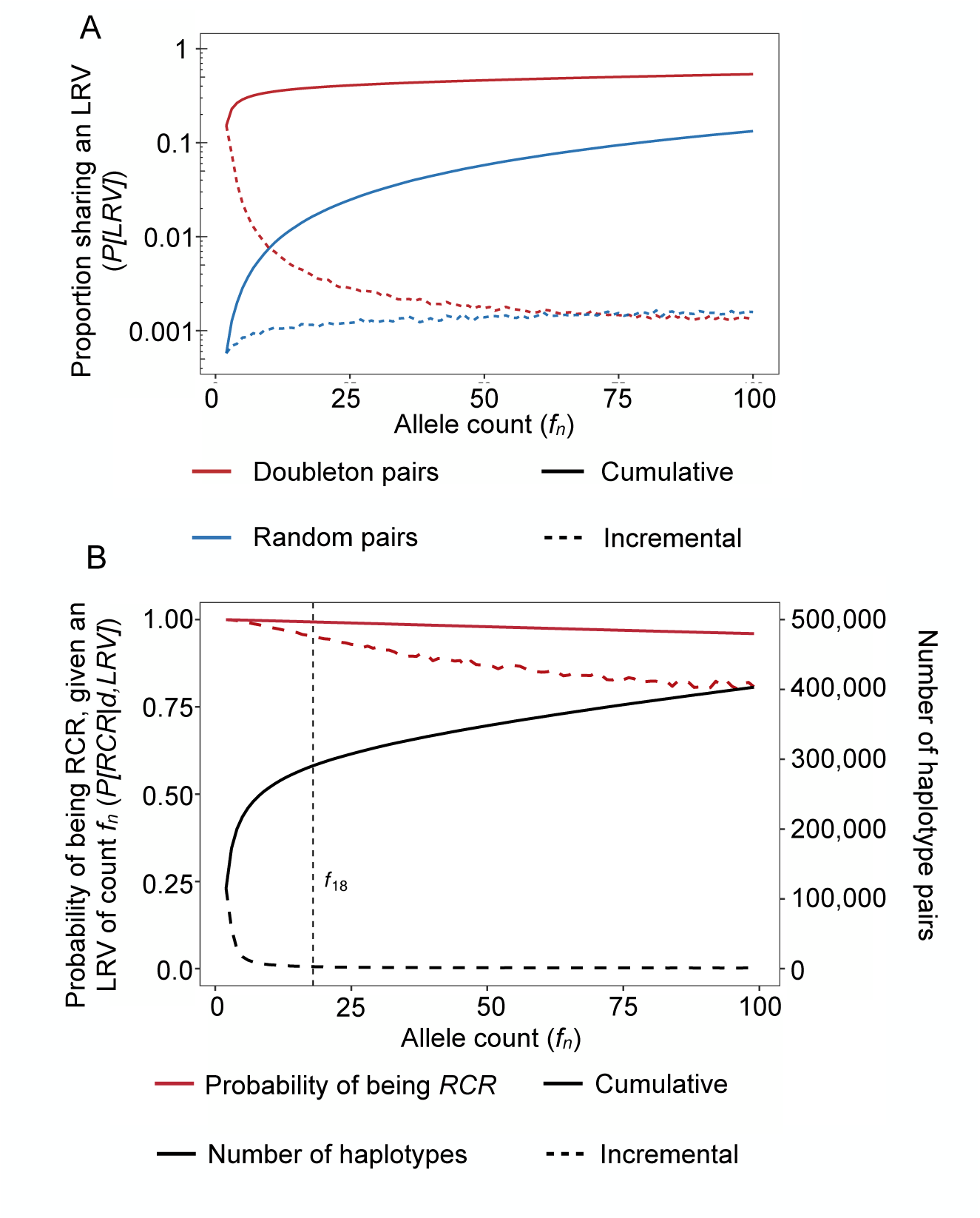
(A) Proportion of doubleton (red) or random (blue) pairs sharing a linked rare variant of count *f_n_*, as a function of the rare variant count. These proportions estimate the corresponding probabilities *P* [*LRV*]. Values are shown for variant allele counts of *n* or less (cumulative; solid lines) and for counts of *n* but not less than *n* (incremental; dashed lines). Both groups were randomly downsampled to 750,000 pairs. (B) Probability that a doubleton with a linked rare variant identifies sequences that are reciprocal closest relatives (*RCR*; red), as a function of the allele count of the rare variant *f_n_*. Solid and dashed lines are as for (A). The number of haplotype pairs with at least one *LRV* of count *f_n_* is also shown (black). The dashed vertical line shows the *f*_18_ threshold used to define the high confidence set of *RCR* doubletons used in this study

#### Identifying *RCR* doubletons

The probability that a doubleton with a linked rare variant has a single origin, *P* [*RCR*|*d, LRV*], was calculated using the preceding results and Eq. (4). We used different threshold values of allele count for the rare variant and, as expected, the lower the frequency of the linked mutation, the greater the probability that the associated doubleton had a single origin, but the smaller the number of doubletons that would be included. These opposing trends are shown in Figure 2B. Based on these results we defined our subset of high confidence *RCR* doubletons as those with linked variants present in 2-18 copies in the dataset (corresponding to a minor allele frequency in the dataset of *<* 0.8%), as the probability that any additional haplotypes added beyond this are *RCR* is *<* 0.95 (Figure 2B; threshold indicated by dashed line). Though doubletons are more likely to have linked singletons than random haplotypes, the presence of a singleton only raises the probability of a doubleton being *RCR* to 0.92, not enough to be included in our high confidence subset. This threshold resulted in a total of 290,704 unique haplotypes being included, carrying 636,938 doubletons, with an overall probability of being *RCR* of 99%, 57% of the total number called and 68% of the estimated number that are *RCR*. By comparison, only *P* [*LRV* |¬*d*] = 2% of the random pair haplotypes shared a linked variant in this range. The vast majority of individuals (1,113 - or 97%) shared at least one *RCR* doubleton with another individual. Each individual shared an *RCR* doubleton with an average of 254 other individuals in the dataset, across 144,542 unique pairs of individuals. As expected, *RCR* doubletons identified in this way have both longer flanking segments than random pair haplotypes (median: 14.3±0.004 vs. 2.9±0.003 kb; p *<* 1*e*−10, Wilcoxon Rank-Sum test) and a higher density of shared *LRVs* up to *f*_18_ (0.3 ± 0.0004 vs. 0 ± 0kb^−1^; p *<* 1*e*−10; Fig. 3A). Moreover, doubletons on longer haplotype segments are more likely to be identified as as *RCR* by our method than those on shorter ones (Fig. 3B).

**Figure 3:**
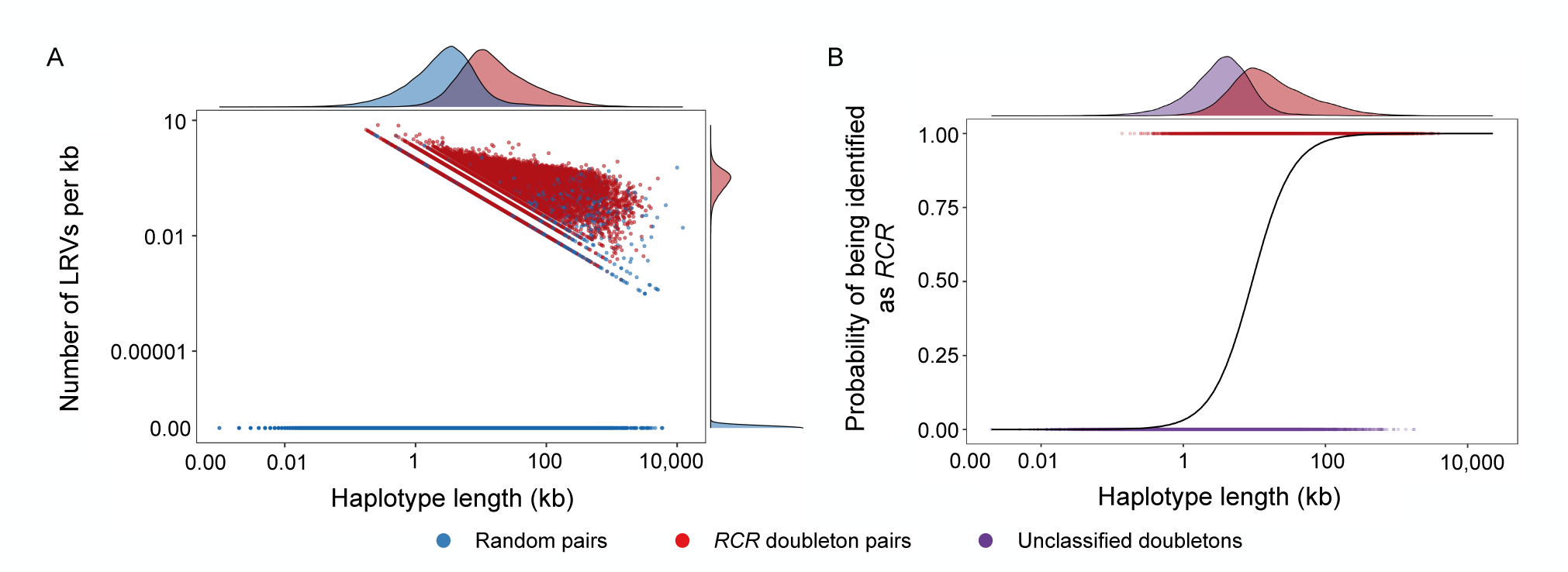
(A) Haplotype length and density of linked rare variants (*LRVs*; ≤*f*_18_) on *RCR* doubleton and random pair haplotypes. Both sets were randomly downsampled to include only 20% of the data to better graphically display differences. Fully 99% of random pair haplotypes had no linked rare variants, constituting the peak at 0. (B) Logistic regression of the probability a doubleton is identified as *RCR* as a function of haplotype length. Marginal plot shows the distribution of haplotype lengths for *RCR* and unclassified doubletons.

#### Assessing the *RCR* classification

As an independent test of the effectiveness of our identification method for high likelihood *RCR* doubletons, we compared the relative frequencies of the different types of mutational change in doubletons with those observed for singletons. For example, C→T singletons are 3.75 times more frequent in the genome than A→C singletons, and if all doubletons have a single origin we would expect C→T doubletons to also be 3.75 times more frequent than the A→C doubletons, whereas if all doubletons are recurrent then C→T doubletons should be 3.75^2^ = 14.1 times more frequent. A 2 x 12 contingency table analysis comparing the frequencies of the 12 different types of mutational change for singletons and our chosen set of *RCR* doubletons shows the two sets are significantly different (G=261.41, *p <* 2.2*e*−10), but the differences are very small, with points very close to the 1:1 line (Fig. 4). By contrast, points for the unclassified doubletons (of which 63% are expected to be *RCR* and 37% recurrent) are closer to the expectation for recurrent mutations.

**Figure 4:**
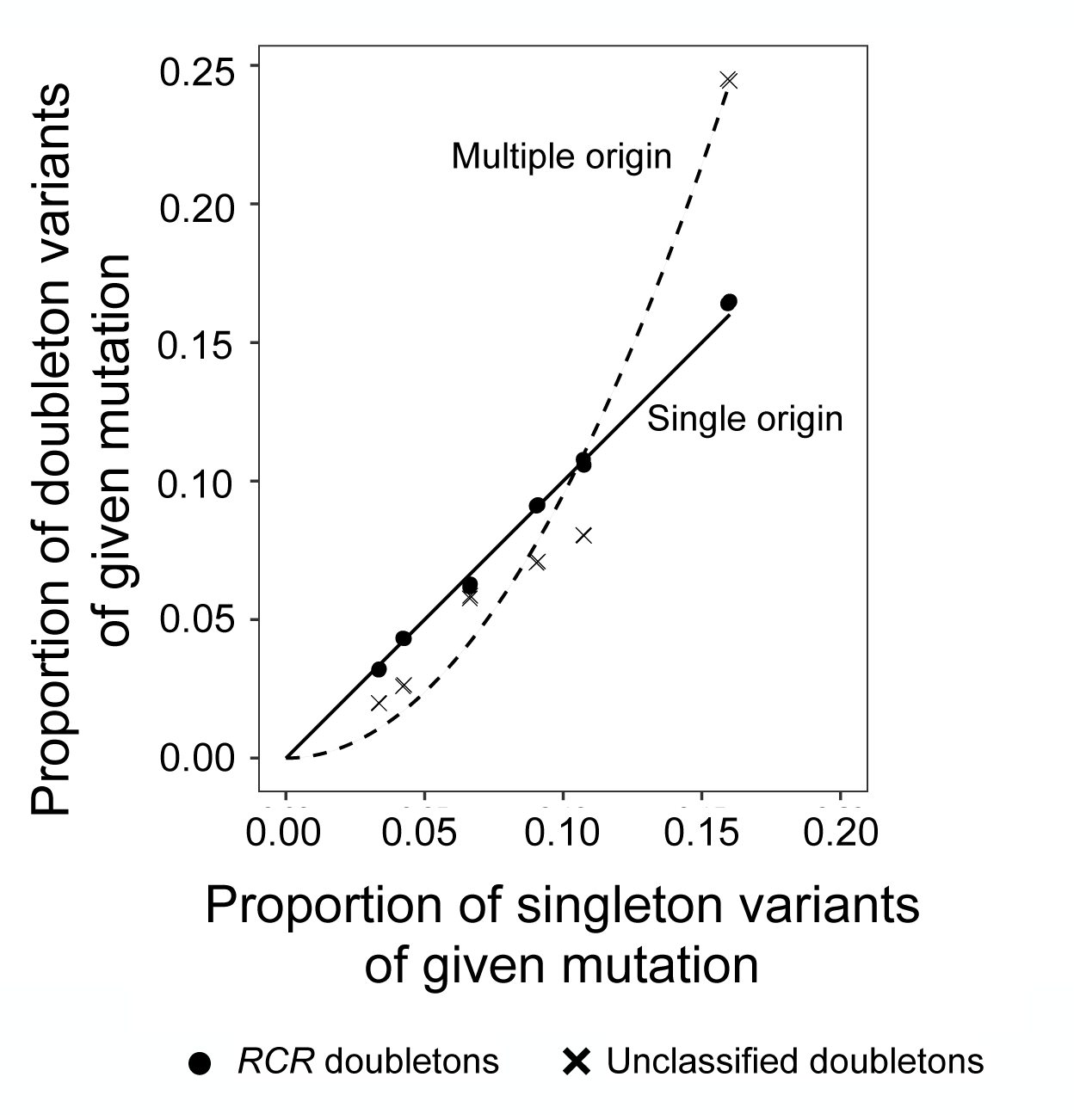
Expected proportions of different classes of doubleton mutations (e.g., A→C) derived from a single origin (i.e., identity-by-descent; 1:1 solid line) and multiple origins (recurrent mutation; dashed line), together with the observed proportions for doubletons classified as *RCR* and those not classified. Expected proportions were calculated from the observed proportions of singleton mutations.

One would expect that, on filtering to include only *RCR* doubletons, much of the ‘noise’ in the data would be removed, enhancing genuine biological signals due to shared ancestry. To test this, differences in patterns of doubleton sharing within and between countries were investigated before and after filtering to include only high-confidence *RCR* doubletons (Fig. 5). It would be expected that sharing would be more pronounced between locations in close geographical proximity and between samples from the same species, when considering only *RCR* doubletons. This was broadly observed, with doubleton sharing more pronounced in location-species clusters (e.g., West African *An. coluzzii*) and reduced between more distant locations (Fig. 5B), with these differences highlighted in Figure 5C.

**Figure 5:**
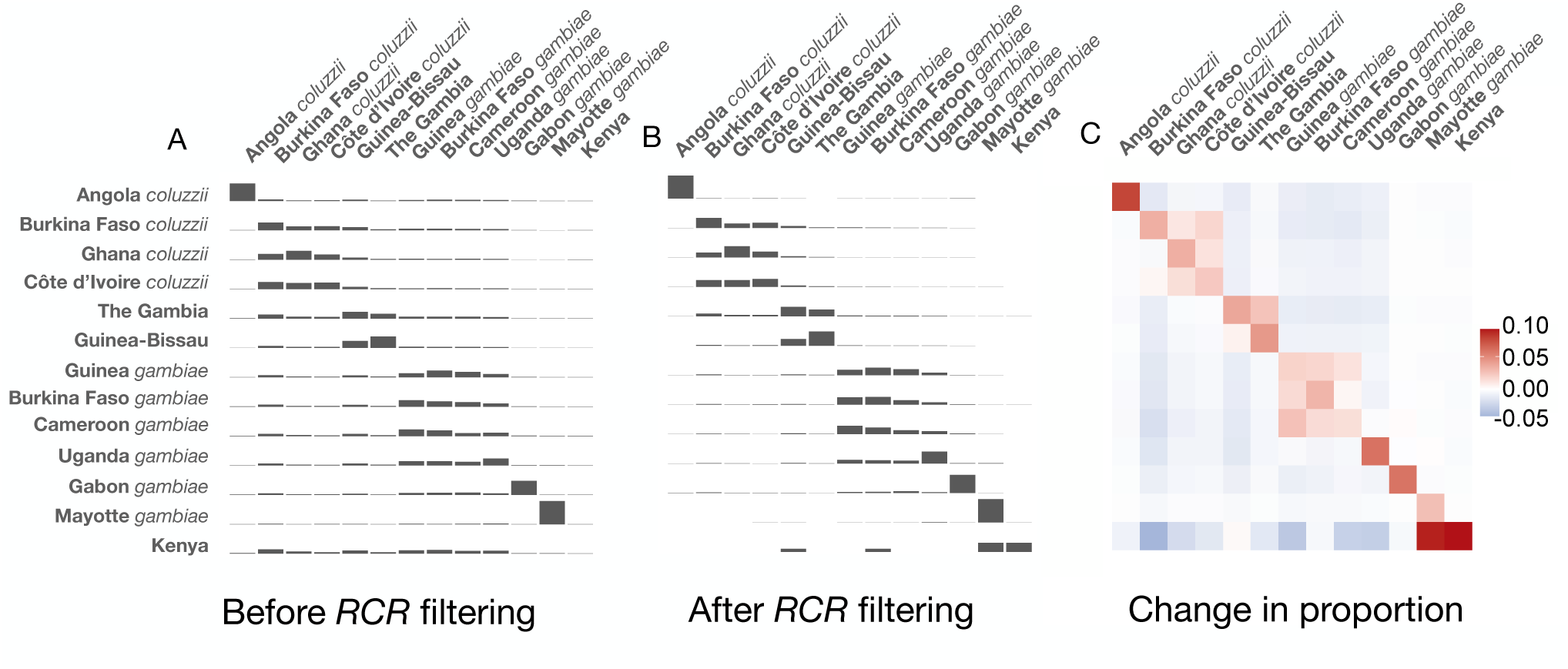
(A) and (B) Doubleton sharing within and between countries, before and after *RCR* filtering. Bar heights are proportional to the number of doubletons shared between individuals of each country combination, and are standardised such that the total height of any given row is fixed. For both panels a random subset of individuals was chosen, equal to the size of the smallest sample (Mayotte, *n* = 24) to remove effects caused by unequal sampling. Groups are loosely ordered in geographical and species clusters. (C) Change in sharing proportions after *RCR* filtering, coloured according to direction of change.

To further test our methodology, we applied it to simulations generated using the coalescent simulator msprime v1.2 (Baumdicker et al., 2022), where information stored in the resulting tree sequences allows one to extract exact information on single versus recurrent origins. Four different demographic scenarios (constant population vs population expansion; panmictic vs island model) were used, to help validate that our methods were not limited to any one singular case. We first examined whether estimates of the proportion of recurrent doubletons obtained from the rates of triallelic singleton sites were comparable to the true values, and found that, for each scenario, the true proportion fell within bootstrapped 95% confidence intervals of the triallelic singleton estimates (Table 2).

**Table 2:** True and estimated recurrent doubleton proportions under different demographic scenarios.

| Demographic scenario*<br>(no. of populations) | Individuals per<br>population | Mutation<br>rate (bp/gen) | Nucleotide<br>diversity ($\pi$) | True recurrent<br>proportion<sup>‡</sup> | Estimated recurrent<br>proportion<sup>§</sup> (95% CI) |
| --- | --- | --- | --- | --- | --- |
| Panmictic, constant (1) | 1e7 | 1.125e-9 | 0.039 | 0.015 | (0.012, 0.016) |
| Panmictic, expansion (1) | 1e8<sup>†</sup> | 3.125e-9 | 0.011 | 0.103 | (0.097, 0.112) |
| Island, constant (5) | 1e6 | 3.125e-9 | 0.054 | 0.020 | (0.018, 0.022) |
| Island, expansion (5) | 1e8<sup>†</sup> | 3.125e-9 | 0.054 | 0.072 | (0.069, 0.075) |

\* 1,000 sampled individuals and a 1Mb genome for each simulation.
<sup>†</sup> Current population size; ancestral population size 1e6, with instantaneous increase to current size 10,000 generations ago.
<sup>‡</sup> From tree sequences
<sup>§</sup> From triallelic singleton data (e.g., Eqs.(2) and (3))

We next determined the association between the inferred probability of a doubleton being *RCR*, given a shared *LRV* of a given frequency, and the true proportion of those doubletons that were *RCR*. If working effectively, one would expect these values to be near identical. In all scenarios there was an excellent correspondence between the two, with our inferred *RCR* probability slightly underestimating the true proportion (Fig. 6).

**Figure 6:**
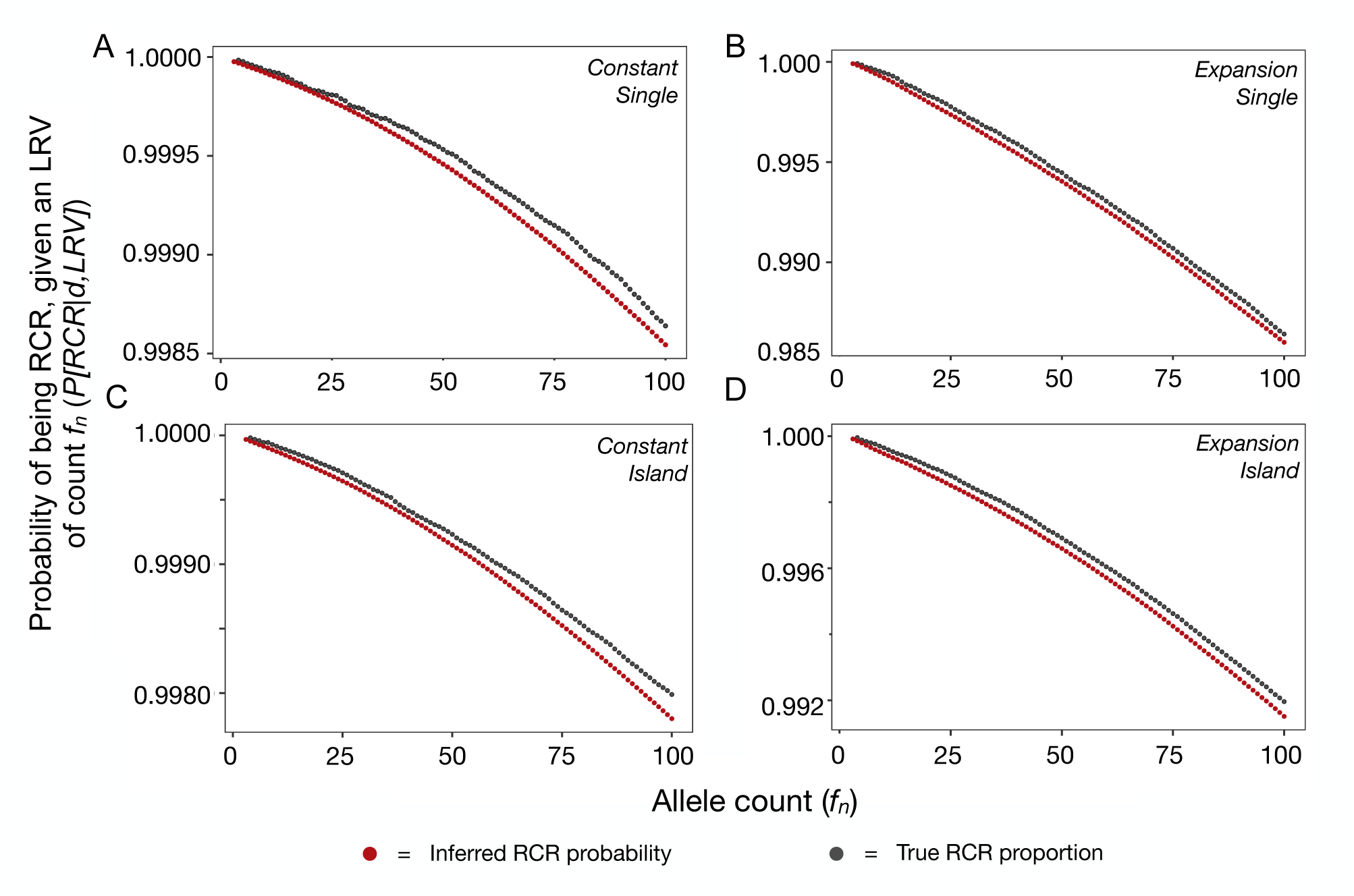
The inferred probability of a doubleton being *RCR* given that it has an *LRV* of count *f_n_* or less, calculated using Eq. (4), and the true proportion of them that are *RCR*, for four simulated demographic scenarios. Values are shown for *LRV* s of frequencies up to *f*_100_. Note difference in y-axis scales.

### Using *RCR* doubletons for population genetic analyses

Having defined a set of high likelihood *RCR* doubletons we now illustrate how they can be used to test population genetic hypotheses involving recombination, selection, and isolation by distance. In many instances these tests involve analysing haplotype lengths, the combined length of the segments from the doubleton out to the first inconsistent homozygote site on each side.

#### Recombination

In many species, including *An. gambiae* s.l., rates of recombination are thought to be lower near the centromere (Choo, 1998), leading to the expectation that doubletons near the centromere will be flanked by longer haplotypes than those further away. Figure 7 shows that haplotypes around both *RCR* doubletons and random pairs are indeed about an order of magnitude longer near the centromere compared to the rest of the chromosome, with this difference decaying over the first 20Mb. The local peaks observed along the chromosome are associated with areas of the genome with no coverage, which increase the distance to the nearest detected inconsistent homozygote site if the haplotype ends in this region. As random pair haplotypes represent the expected distance to an inconsistent homozygote site from a given position in the genome, these should give a rough indicator of the length of the overestimate generated by our approach to haplotype detection. The difference between these and the *RCR* haplotypes, plotted in black, should therefore provide an estimate of the true length of *RCR* haplotypes in a given region.

**Figure 7:**
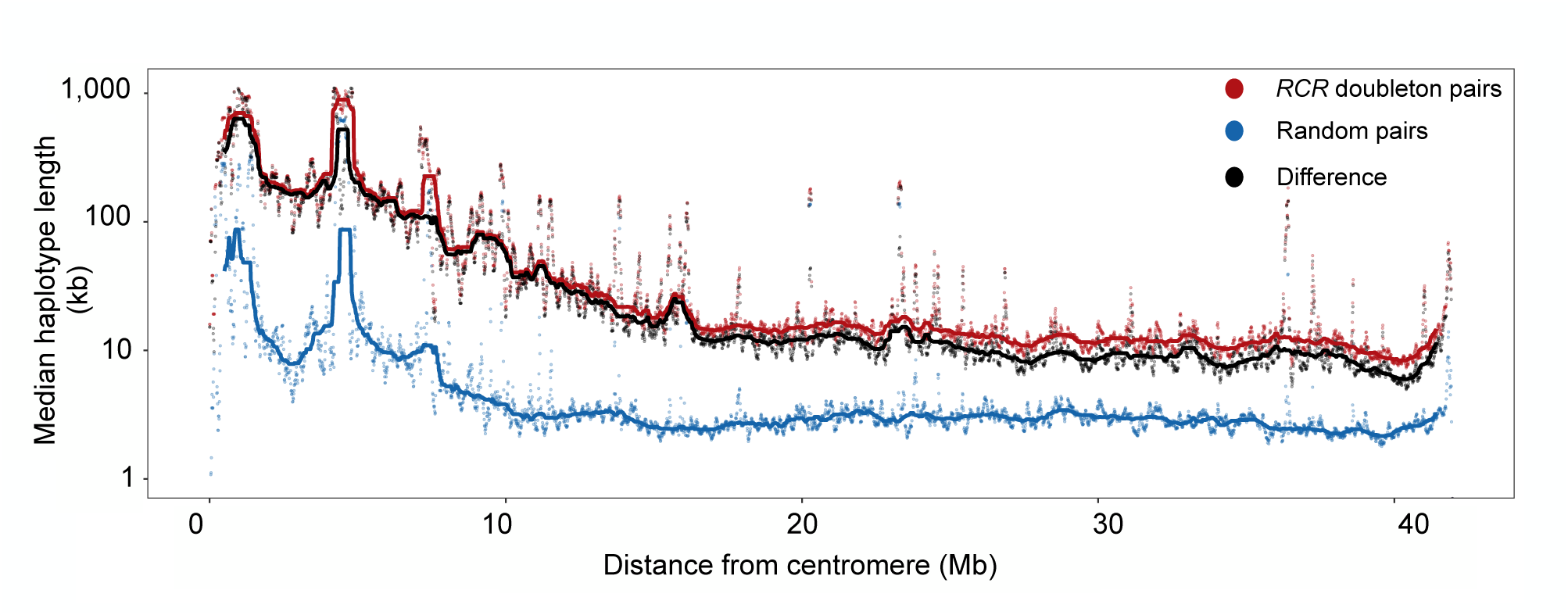
Effects of chromosome position on the length of haplotypes flanking *RCR* doubletons. Median lengths of *RCR* and random pair haplotypes are shown along chromosome arm 3L, and the difference between the two groups at each position, highlighting elevated haplotype length close to the centromere, a likely effect of reduced recombination. Position is defined by the midpoint of the haplotype span, with each section of the chromosome binned into 10kb regions (points) and then smoothed via a rolling median spanning 100 bins (lines).

#### Selection

At least some of the doubleton mutations are likely to reduce fitness and be selected against, and the stronger the selection against the mutation, the more recent the mutation is expected to be, and therefore the longer the haplotype (Maruyama, 1974; Kiezun et al., 2013; Mathieson and McVean, 2014). To test whether the data fit these expectations we first compared the length of haplotypes associated with doubletons occurring at zero-fold and four-fold degenerate sites, as mutations in the former are more likely to be deleterious than those in the latter (Li et al., 1985). As expected, haplotypes flanking zero-fold doubletons are significantly longer than those flanking four-fold doubletons (median 20.5 ± 0.33 vs. 16.0 ± 0.30kb, *p <* 2.2*e*−10, Wilcoxon Rank-Sum test), though there is considerable overlap in the distributions (Table 3).

**Table 3:** Haplotypes associated with doubletons at positions of different codon degeneracy or nucleotide conservation.

| | Number of<br>haplotypes | Mean length<br>± SE (kb) | Median length<br>± SE (kb) | Length quartiles<br>(lower; upper - kb) |
| --- | --- | --- | --- | --- |
| *Codon Degeneracy* |  |  |  |  |
| Four-fold | 5,737 | $46.7 \pm 1.52$ | $16.0 \pm 0.30$ | 7.6; 40.1 |
| Zero-fold | 9,631 | $62.6 \pm 1.49$ | $20.5 \pm 0.33$ | 9.0; 55.4 |
| *Nucleotide Conservation* |  |  |  |  |
| Non-conserved site* | 2,893 | $52.8 \pm 2.46$ | $16.6 \pm 0.48$ | 7.7; 43.8 |
| Conserved site<sup>†</sup> | 16,618 | $55.3 \pm 1.08$ | $17.7 \pm 0.24$ | 8.1; 46.9 |
| Conserved region<sup>‡</sup> | 274 | $80.2 \pm 13.6$ | $21.4 \pm 2.41$ | 9.8; 54.6 |

\* Not conserved across 21 species of *Anopheles*
<sup>†</sup> Conserved across 21 species of *Anopheles*.
<sup>‡</sup> Inside of a region of at least 18 contiguous conserved sites.

Nucleotide bases that are conserved across species are also more likely to be under stronger purifying selection than sites that are not conserved, and therefore we used an alignment of sequences across 21 species of *Anopheles* mosquitoes and categorised doubletons as being at (i) non-conserved sites; (ii) completely conserved sites across the 21 species; and (iii) completely conserved sites that are also inside a region of at least 18bp that is also completely conserved. Again, haplotype lengths showed small but statistically significant differences in the expected direction (median 16.6 ± 0.48 vs. 17.7 ± 0.24 vs. 21.4 ± 2.41kb, all *p <* 0.05, Wilcoxon Rank-Sum tests (Table 3).

#### Isolation by distance

The sequences in the Ag1000 Phase 2 dataset are from 13 countries across a broad swath of subSaharan Africa, spanning distances much larger than the typical dispersal distance of an individual mosquito, and therefore individuals collected near to each other are expected to be more closely related than those collected further apart. This higher relatedness should be manifest both in a higher probability of them sharing an *RCR* doubleton and in the haplotypes flanking those doubletons being longer. The data support these predictions. First, 48% of *RCR* doubletons are shared between individuals from the same country, compared to 21% of the unclassified doubletons, and a random null expectation of 11% (*G* = 175, 560*, p <* 2.2*e*−10; Fig. 8A). Second, *RCR* doubletons from individuals from the same country have significantly longer flanking haplotypes than those from different countries (Fig. 8B). This difference is found for doubletons in both *An. coluzzii* (median; 25.2 ± 0.23 vs. 15.2 ± 0.14kb; *p <* 1*e*−10, Wilcoxon Rank-Sum test) and *An. gambiae* s.s. (median; 19.7 ± 0.12 vs. 10.7 ± 0.04kb; *p <* 1*e*−10, Wilcoxon Rank-Sum test). Finally, considering only *RCR* doubletons from individuals from different countries, there is a negative correlation in both species between the geographical distance separating the sampling locations and haplotype length (Fig. 8C). Here, the haplotype length considered is the total length of all unique haplotypes shared between each pair of individuals, to account for some pairs of individuals who share multiple *RCR* haplotypes with each other (38% of those between countries), often in close genomic proximity. Interestingly, *An. coluzzii* has both longer haplotypes than *An. gambiae* s.s. within countries (*p <* 1*e*−10, Wilcoxon Rank-Sum test), and a steeper decline in haplotype length with geographical distance between countries (*p* = 0.021). Random pair haplotype lengths also decrease with geographical distance, as expected with isolation by distance, though with a shallower slope.

**Figure 8:**
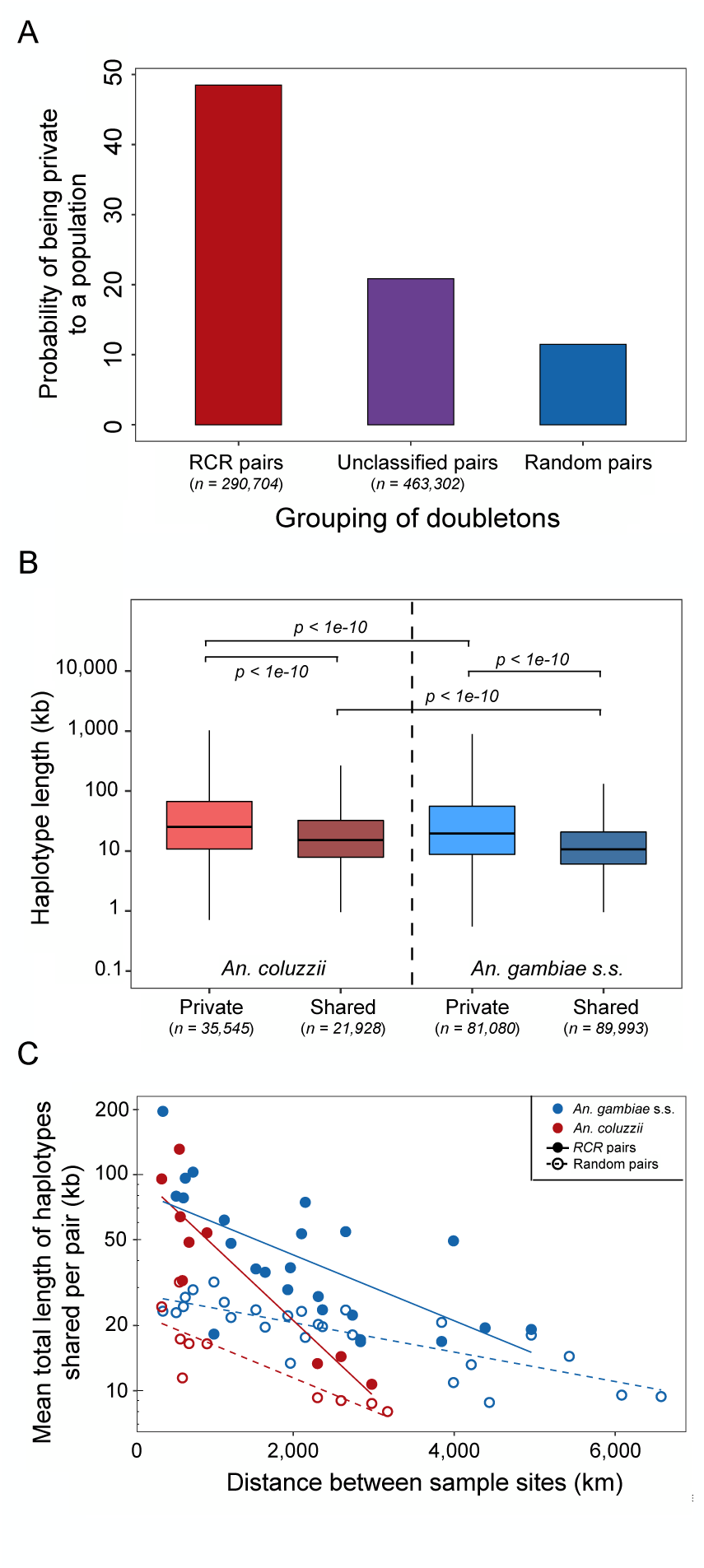
Relationship between genetic relatedness inferred from *RCR* haplotypes and geographical distance. (A) Probability of a doubleton being found between individuals within a country (private) for *RCR* and unclassified doubletons. Also shown is the null expectation if doubletons were randomly distributed among pairs of individuals. Error bars were too small to be displayed on the graph. (B) Distribution of haplotype lengths for doubletons shared within and between countries, for *An. coluzzii* and *An. gambiae* s.s. Significance of pairwise Wilcoxon Rank-Sum tests is displayed. (C) Mean total length of shared *RCR* haplotypes per pair of individuals, between countries, as a function of the geographical distance between sites for *An. coluzzii* (red solid points and line) and *An. gambiae* s.s. (blue); the slopes of Ln[length] on distance (in km) are −0.00081±0.00014, *R*^2^ = 0.808*, p* = 6.1*e*−4 and −0.00035±0.00008, *R*^2^ = 0.423, *p* = 4.6*e*−4, respectively. Also shown are the mean lengths of shared haplotypes for the random pairs of individuals (open circles and dashed lines; slopes 0.00035 ± 0.00008, *R*^2^ = 0.679*, p* = 2.1*e*−3 and 0.00015 ± 0.00002, *R*^2^ = 0.631*, p* = 2.7*e*−7 for *An. coluzzii* and *An. gambiae* s.s., respectively).

## Discussion

### Identifying *RCR* doubletons

The usefulness of a molecular marker for population genetic inference is in large part a function of how likely it is to have a single origin, and therefore identifies individuals that are genealogically related. Rare variants are of interest because they are more likely than common variants to have a single origin and to have arisen relatively recently, and may therefore allow for finer discrimination of populations (e.g., Gravel et al., 2011; O’Connor et al., 2015). In this paper we have focused on doubletons, the rarest of rare variants that can be informative about relatedness between individuals, and used them to analyse published data from *An. gambiae* s.l. mosquitoes. While it is commonly assumed that their rarity ensures derivation from a single common ancestor, this may not be the case and could confound analyses if not properly accounted for. We first devised an approach for estimating the frequency of doubletons that have single origins, based on the frequency of triallelic singleton sites. We then developed a method to identify a subset of doubletons with a high likelihood (ca. 99%) of being single origin, based on the presence of a linked rare variant. Applying these methods to published genomic data from *An. gambiae* s.l. mosquitoes, we estimated that 5-25% of doubletons are due to recurrent events, depending on the type of mutation, with an overall estimate across all doubletons of ∼16%. We then identified a subset of ∼637, 000 doubletons (out of a total of ∼1, 100, 000) that are highly likely to have single origin, which define ∼291, 000 unique haplotype pairs. As a test of our methodology, we then checked (1) that the mutational spectrum of these doubletons matches that of singleton mutations; (2) an analysis of doubleton sharing within and between countries shows enhanced within-country sharing, consistent with the removal of ‘noise’ due to recurrent mutations; and (3) coalescent simulations under a variety of demographic scenarios showed a close correspondence between predicted and true values.

A number of factors are expected to affect the proportion of doubletons that are single origin in a population genomic dataset. A recurrent doubleton occurs if the same mutation arises on two different external branches of the genealogy at a site (and on no internal branch). Under the standard neutral model of constant population size, random mating and no selection, the average or expected total length of the external branches of a genealogy is 4Ne (where Ne is the effective population size) and is independent of sample size (Fu and Li, 1993). Therefore, the probability of there being two mutations on external branches will be higher when the effective population size is larger and/or the mutation rate is higher, but will be (largely) independent of sample size (see Wakeley et al. (2023) for a more formal treatment). However, if the population size has not been constant, then sample size can have an effect: increasing the sample size reduces the average length of external branches (i.e., their first coalescent event occurs more recently than if the sample size was smaller), and the rate of those coalescent events will be more influenced by recent population size. Thus, in populations that have been expanding (like *An. gambiae* s.l.; (Khatri and Burt, 2019; The Anopheles gambiae 1000 Genomes Consortium, 2020)) the total length of external branches is expected to increase with sample size, and the proportion of recurrent doubletons as well (Wakeley et al., 2023). Note too that a high frequency of sequencing errors could also increase the proportion of (apparent) doubletons that are recurrent. For this study we followed the relatively stringent quality thresholds employed by the Ag1000G Project (2020). To the extent that sequencing errors can be considered an additional source of mutations on external branches, and would lead to triallelic singleton sites as well as doubleton sites, our estimate for the number of single origin doubletons should be largely unaffected.

To identify a subset of doubletons that are highly likely to have a single origin, we developed a method using the formalism of Bayes’ Theorem and the different proportions of doubleton haplotypes and matched random pair haplotypes that have a linked rare variant (*LRV*). We defined ‘linked’ based on the haplotype detection approach of Mathieson and McVean (2014), and varied the definition of ‘rare’ to arrive at a desired confidence level. *RCR* doubletons are expected to differ from matched random pairs of individuals both in the length of their flanking haplotypes and in the density of *LRVs* found on them, and, by using the presence of an *LRV* anywhere in the haplotype, our method indirectly combines both features to identify high likelihood *RCR* doubletons.

The only previous method we are aware of for distinguishing between rare variants with single or multiple origins is that proposed by Johnson et al. (2022), which is based on haplotype length alone. Their method also differs in that it requires a demographic model of the study population - specifically assuming a single, exponentially growing population - whereas ours does not appear to make any assumption about demographic history. It does assume that (1) doubletons are derived alleles with either 1 or 2 origins, with no back mutations; and (2) that triallelic singletons are both derived alleles from the common type, again with no back mutations. It is possible that the method may break down if there are many sites with many (*>>* 2) mutations in the history of the species (e.g., RNA viruses or some localised hypermutability). Further simulations would be needed to assess this possibility. Under such a hypermutable scenario, distances to the nearest incompatible homozygous sites might under-estimate (rather than over-estimate) the distance to a recombination site, which may reduce the power to detect *RCR* doubletons.

The median length of our high confidence *RCR* haplotypes was 14.3kb. Using a genome-wide average recombination rate of 1.8 cM/Mb (The Anopheles gambiae 1000 Genomes Consortium, 2017), this translates to about 0.03cM. These values are much shorter than were found in Mathieson and McVean’s (2014) analysis of doubleton haplotypes in human data (600kb and 0.7cM), with roughly comparable sample size and recombination rates. The much smaller values in mosquitoes are consistent with haplotypes being much older (i.e., more time back to the common ancestor), which in turn is consistent with mosquitoes having a much larger population size. Note too that 0.03cM is 100-fold shorter than the limit of confident detection of IBD segments in standard human analyses (e.g., Browning and Browning, 2013) (though see Shetty et al. (2020) who use *LRV* s to improve confidence for shorter segments). This difference is presumably due to a combination of the methodology used and the higher density of polymorphic sites in mosquitoes. One way to recover longer *RCR* haplotypes, with more recent common ancestors, would be to increase the sample size as this would increase the relatedness of the closest relatives. In the standard neutral model, if the sample size is doubled, then the average length of external branches is halved, and so is the distance to the nearest relative (Fu and Li, 1993). In *An. gambiae* s.l., where the population has likely been expanding, the effect of doubling the sample size will not be as large, but still one can probably expect more than twice as many detectable haplotypes for downstream analyses.

### Analyses using *RCR* doubletons

Having defined a high confidence subset of single origin doubletons, we then used them to test a series of population genetic hypotheses. We find that *RCR* doubletons near the centromere are associated with longer haplotypes, as expected if recombination rates there are reduced. Second, *RCR* doubletons at sites likely to be under purifying selection are associated with longer haplotypes, consistent with them being more recent, a difference also seen in human data (Kiezun et al., 2013; Mathieson and McVean, 2014). This relationship could be further explored by, for example, using a continuous variable like Genomic Evolutionary Rate Profiling (GERP) scores instead of the simple binary variable conserved vs non-conserved (Cooper et al., 2005; Davydov et al., 2010). And third, the data are consistent with an isolation by distance model: *RCR* doubletons are more likely than random pairs to be derived from two individuals in the same country; *RCR* doubleton haplotypes in the same country are longer than those from different countries; and for those shared between countries, the combined lengths of their haplotypes decreases with geographical distance between the locations.

We also found differences in haplotype lengths between the two *An. gambiae* s.l. species. Within countries, haplotypes are longer in *An. coluzzii*, consistent with a smaller average local population size. Haplotypes also shorten more rapidly in *An. coluzzii* with geographical distance, indicating stronger isolation by distance. Previous work has shown that *F*_ST_ between countries increases more rapidly with geographical distance in *An. coluzzii* than in *An. gambiae* s.s. (The Anopheles gambiae 1000 Genomes Consortium, 2020), an analysis based on a completely different aspect of the same population genomic data (i.e., differences in allele frequencies, independent of linkage). While the relationship between *F*_ST_ and geographical distance is expected to depend only on the product of population density and dispersal distances, these parameters are in principle separable in haplotype-based analyses (Barton et al., 2013). Our analyses appear to suggest that *An. coluzzii* not only has a lower average local population size, but also a shorter average dispersal distance. Shorter dispersal distances for *An. coluzzii* could make sense ecologically, as they are thought to prefer permanent water sources, as opposed to *An. gambiae* s.s. which prefer more ephemeral sources (Kamdem et al., 2012; Akpodiete et al., 2019). Moreover, there is a greater evidence for *An. coluzzii* aestivating in the Sahelian dry season and therefore may not be so reliant on dispersal to persist (Dao et al., 2014). On the other hand, of the two species, *An. coluzzii* has thus far been found more often at high altitude during airborne sampling, where it is thought upper atmospheric currents may facilitate long range dispersal (Huestis et al., 2019). In principle, differences in mutation rates or recombination rates may also contribute to haplotype length differences between species.

### Future developments

In principle, it is possible that our methods could be modified to recover a greater proportion of high confidence *RCR* doubletons. Our analysis of triallelic data showed that different classes of doubletons have different prior probabilities of having a single origin, but in the subsequent analysis incorporating linked rare variants, for simplicity, we only used the overall prior probability and derived a single threshold frequency. We could instead have used mutation-specific prior probabilities of being single origin, and calculated separate thresholds for each. One could also go beyond this to incorporate differences in mutation rate due to trinucleotide or other local sequence context (Hwang and Green, 2004; Harris and Pritchard, 2017; Mathieson and Reich, 2017; DeWitt et al., 2021). A further extension could also take into account the number of linked rare variants (e.g., if the current threshold is *f*_18_, maybe having 4 *f*_25_’s would also be sufficient). Our method identified 68% of the estimated number of *RCR* doubletons based on the analysis of triallelic sites. It is possible this is an under-estimate, if the expected number was itself an over-estimate, and it is possible that the modifications above might increase the number identified. However, there is presumably an upper limit to the fraction of single origin doubletons that are detectable by testing for linked rare variants - for example, if one copy has been transferred to a different genetic background by a gene conversion event with no other rare variant in the conversion tract, then it will be missed. Such short haplotypes would in any case seem difficult to date (i.e., would contain little information on the degree of relatedness), and therefore a failure to identify them may not be such a loss.

The analysis could also be extended in different ways. First, to avoid uncertainties associated with statistically phasing the sequence data prior to analysis, we used unphased data (though the method of analysis does mean that, for single origin doubletons, the doubleton and linked rare variant are highly likely to be on the same chromosome). Some studies have found that errors in inference introduced by incorrect imputation can be greater than that introduced by using unphased and therefore less accurately identified haplotypes (Janzen and Miró Pina, 2022). If robust phasing data is available the method could be modified to incorporate it. Second, one could go beyond doubletons to mutations present in 3, 4, or more individuals. One approach may be to do the same analysis for all possible doubletons in a tripleton, and look for congruence in sharing of *LRVs* — if all pairs of a tripleton are judged as single origin, then that would suggest that the tripleton is a clade. Interestingly, for a fixed mutation rate, the distribution of the number of origins of a rare variant appears to rapidly become independent of the allele count once it is above a relatively small number (Wakeley et al., 2023).

The other main extension is in parameter estimation. We have shown that the length of the haplotype on which an *RCR* doubleton resides varies in expected ways with chromosomal position, with whether or not the doubleton is likely to be under purifying selection, and with geography. The data thus contains information on recombination rates, fitness differences, and movement, and a worthwhile next step, beyond the scope of this paper, would be to develop the appropriate inferential methods for using these data to estimate the parameters of a model. Such estimates will be especially useful for some purposes because they should reflect relatively recent parameter values. In the study of human population genetics, IBD haplotypes have typically been defined based on more common variants, excluding rare variants (e.g., Browning and Browning, 2013; but see Shetty et al. (2020)), and have been very useful for demographic inference in humans (e.g., Palamara et al., 2012; Barton et al., 2013; Ralph and Coop, 2013; Baharian et al., 2016; Ringbauer et al., 2017). The challenge is that defining IBD haplotypes will be more difficult when the ratio of recombination rate to mutation rate is high, and while in humans this ratio is about 1, in *An. gambiae* s.l. it is about 10 (Nelson et al., 2021). Using rare variants to define haplotypes, as demonstrated here, may be a useful way forwards. Given how much harm is caused by this species (World Health Organisation, 2020), and that population genetic interventions including using gene drive are being actively called for (African Union, 2018; World Health Organisation, 2021) and developed (e.g., Hammond et al., 2021; Hoermann et al., 2022; Carballar-Lejarazú et al., 2023), further investigation of appropriate methods to extract as much information as possible from population genomic data is much to be desired.

## Materials and Methods

### Data acquisition

Genomic data was acquired from the Phase 2 AR1 data release of the *Anopheles gambiae* 1000 Genomes project, in VCF format, for chromosome arm 3L (https://www.malariagen.net/data/ag1000g-phase-2-ar1). This release consisted of 1,142 wild-caught individuals from 13 countries across subSaharan Africa, belonging to the sister species *An. gambiae* s.s. and *An. coluzzii*, who comprise two of the primary species in the *Anopheles gambiae* species complex (*An. gambiae* s.l.). As described in the main text, unphased reads were used due to concerns over phasing of rare variants. In addition to variant calls, sample metadata were also accessed from the data release.

### Data filtering

Initial filtering for quality was performed using the pre-prepared PASS filter from Ag1000G. This removed any SNP that was not classified as ‘accessible’, removing sites subject to poor sequencing quality, allelic dropout, poor alignment, and with problematic repeats and structural variation. Furthermore, it provided additional quality control to account for allelic position on reads, mapping of repeats, strand bias, and coverage relative to depth. For more detail on the exact filters used, see Supplementary Information 3.4 & 3.5 in The Anopheles gambiae 1000 Genomes Consortium (2017). Following quality filtering, SNPs were further filtered to remove any site where there was missing data – that is, every sampled individual had two SNP calls at that position. For haplotype detection, for both doubleton and random pairs of individuals, SNPs were finally filtered to include only those at biallelic sites.

### Haplotype detection

Haplotype detection was performing using modified versions of the scripts from Mathieson and McVean (2014), found at https://github.com/mathii/f2. In short, VCFtools (Danecek et al., 2011) was used to identify sites with doubletons and the individuals carrying them; subsequently, the combined homologous chromosomes of each pair were parsed in both directions from the focal doubleton until a site where the two individuals had inconsistent homozygote genotypes was encountered, which can be inferred to be an estimate of the end of the haplotype as at this position the individuals do not share identity-by-state (and thus identity-by-descent). Any doubletons which were found in a single homozygous individual, and those haplotypes reaching either end of the chromosome arm, were discarded. About 5% of haplotypes had directly adjacent start/end points with another haplotype and it is possible these do not represent separate haplotypes, but we did not investigate them further.

For the random pair haplotypes, two individuals were randomly sampled at each position where a doubleton was identified, but that did not carry the doubleton themselves. Shared haplotypes were then identified in the same manner as for doubleton pairs. For both groups, lengths were identified as the difference between the positions of the two inconsistent homozygote genotypes.

### Allele counting

To determine what linked variants lay on each shared haplotype, VCFtools was used to identify the position and individuals carrying variants of counts from 3 to 100 in the sample. For each haplotype pair, the number of variants of each count shared by both individuals, within the bounds of the shared haplotypes, was recorded. Density of linked rare variants was calculated using the number of variants of counts 2-18 (our threshold value) shared on each haplotype, divided by haplotype length. Counting of singletons, doubletons, and triallelic singletons of each specific base change was performed using scikit-allel (Miles et al., 2021).

### Confidence intervals

Confidence intervals for the proportion of recurrent doubletons estimated from triallelic singleton frequencies were calculated by bootstrapping. Multinomial sampling was performed with event probabilities *X*, *Y*, *Z*, and 1 − *X* − *Y* − *Z*, where *X*, *Y*, and *Z* correspond to the frequencies of the three possible triallelic singletons from a given base in the population (thus 1 − *X* − *Y* − *Z* to the frequency of sites for which there are none), and the number of trials equal to the number of sites in the genome with that base as the major allele. This process was performed for each base (i.e. 4 times in total), and used to obtain estimates of the expected number of recurrent doubletons given these observed frequencies. Subsequently, 10, 000 replicates were produced to give an expected distribution, from which 95% confidence intervals were determined.

### Simulations

Simulations were performed using the coalescent simulator msprime v1.2 (Baumdicker et al., 2022). Four different demographic scenarios were considered: (1) a single population of constant size N=10,000,000; (2) a single population initially of size N=1,000,000 that instantaneously expanded 100-fold to N=100,000,000 10,000 generations ago; (3) an island model with 5 populations each of constant size N=1,000,000 and symmetrical per-generation migration rates of 0.01 between each population; and (4) an island model with 5 populations each initially of size N=1,000,000 that expanded 100-fold to N=100,000,000 10,000 generations ago, with the same migration rates (Table 2). Parameters were chosen such as to give a range of conditions over which to test. For each simulation, the ancestry of 1,000 diploid individuals (200 per population for the island models) was simulated using Hudson’s coalescent, with a recombination rate of 1.8e-8 M/bp, based on genomewide average estimates for *An. gambiae*(The Anopheles gambiae 1000 Genomes Consortium, 2017). Simulations were run until complete coalescence across all sites in the genome. Due to computational constraints on the available high performance cluster, each simulation covered a genome of 1Mb. For each scenario, 70 replicates were completed, and the results pooled to emulate a 70Mb genome. Only ∼1% of observed doubleton haplotypes reached the ends of their simulated chromosomes meaning that this partitioning is unlikely to have introduced undesired artefacts.

For each scenario, tree sequences were overlaid with mutations at rates that led to realistic levels of nucleotide diversity (*π*) (Table 2) and the position and number of true recurrent doubletons were identified using information stored in the tree sequences. To mimic the processes employed on the empirical data, these mutated tree sequences were converted into VCFs using tskit v0.5.8(Kelleher et al., 2016; Ralph et al., 2020; Wong et al., 2024), allowing haplotype detection and LRV counting to be performed in an identical manner. Triallelic singleton based estimates of recurrent doubleton rates were compared to true rates by performing bootstrapping as described above, with values averaged across all simulations for a given scenario. For each simulation, doubleton haplotypes were identified, and their probability of being *RCR*, given shared *LRV* s with allele counts up to 100, calculated. The true proportion of doubleton haplotypes that were *RCR* was then determined for those sharing an *LRV* of each of these counts by cross-referencing with the true recurrent sites identified directly from the tree sequences.

### Site classifications

Positional filters to determine the fold degeneracy of sites in the genome were generated using scripts supplied by Alistair Miles (unpublished), and then applied to doubleton sites. Filters to determine the conservation of sites (non-conserved, conserved site, conserved region) were supplied by Samantha O’Loughlin (personal communication). Conserved sites were identified via the alignment of 21 species within the *Anopheles* genus, with conserved regions referring to those sites which lie in contiguous segments of conserved sites 18bp or more in length (O’Loughlin et al., 2021). For both degeneracy and conservation, where there were multiple doubletons on a single haplotype, each doubleton was considered its own entity.

### Isolation by distance

Geographical distances between sample sites were calculated using associated latitude-longitude coordinates from Ag1000G sample metadata. Where collections had been performed at multiple sites in an area, an average position of these coordinates was used. For comparisons between different countries, average total haplotype lengths were calculated on a per unique pair basis – that is, for every pair of individuals that shared a haplotype, the length of all of the haplotypes they shared between them were combined, to give a total length of shared haplotype. Pairs of countries for which there were *<*10 unique pairings of individuals with shared haplotypes were excluded from further analysis. Pairwise comparisons of haplotype lengths were computed using pairwise Wilcoxon RankSum tests with a Holm-Bonferroni correction. Difference in slope was tested via the interaction term of a linear regression model.

## Acknowledgements

We would like to thank Matteo Fumagalli, John Connolly, and Ace North for their comments on this manuscript, Martin Donnelly and Alfried Vogler for useful discussions, and Joshua Schraiber and an anonymous reviewer for insightful feedback on an earlier version of this paper. This work was supported by grants from the Gates Foundation and the Open Philanthropy Project.

## Data availability

Genomic data used was acquired from the publicly available Phase 2 AR1 data release of the *Anopheles gambiae* 1000 Genome project (which can now be accessed via https://malariagen.github.io/vector-data/landing-page.html). Code used is available at https://github.com/JoshJRey/doubRCR.

