## Supplementary Information for "Identifying single origin rare variants in population genomic data"

### Derivation of Eq. (1) in Main text

Equation 1 from the main text seeks to estimate the probability of any given doubleton being recurrent, depending on the nucleotides concerned. It does this by the use of relative rates of triallelic singleton sites in the genome (those sites with two alternative singleton mutations), with the logic being that these are sites that, under the most likely scenario, have observably been subject to exactly two independent mutations, as one would expect in the case of a recurrent doubleton. If sites of a higher frequency were considered, one would not be able to exactly determine how many recurrent mutations had occurred.

Following the example in the main text, we will assume the ancestral base at a particular site is A, and define $P[A \to C]$, $P[A \to G]$, and $P[A \to T]$ as the probabilities that a mutation of an A on an external branch of the genealogy results in a C, G, or T, respectively, where the three probabilities sum to 1. Conditional on a site being ancestrally A and that there have been two mutations on external branches, the probability that one of the mutations is to a C and the other to a G is then

$$P[A \to C, G] = 2P[A \to C] \times P[A \to G] \tag{SI-1}$$

and similarly for the other combinations. The factor of 2 is needed because there are two possible ways this arrangement could arise: the 1st mutation could be a C and the second a G, or vice versa. The probability that both mutations were to a C (i.e., it is a recurrent doubleton) is

$$P[A \to C, C] = P[A \to C]^2 \tag{SI-2}$$

There is no factor of 2 because there is only one way this arrangement could arise. If we (a) convert to a fraction and then multiply top and bottom by $P[A \to G]$, $P[A \to T]$, and two factors of 2, (b) collect terms appropriately; and (c) use the definition in (SI-1), we get:

$$P[A \to C, C] = P[A \to C] \times P[A \to C]$$

$$= P[A \to C] \times P[A \to C] \times \frac{P[A \to G]}{P[A \to G]} \times \frac{P[A \to T]}{P[A \to T]} \times \frac{2}{2} \times \frac{2}{2}$$

$$= \frac{(2P[A \to C]P[A \to G])(2P[A \to C]P[A \to T])}{2(2P[A \to G]P[A \to T])}$$

$$= \frac{P[A \to C, G]\ P[A \to C, T]}{2P[A \to G, T]} \tag{SI-3}$$

which is Equation (1) in the main text.

### Estimating the proportion of *RCR* doubleton sites: an example calculation

Here, we provide an example of our approach to estimating the proportion of doubleton sites in our dataset that should be *RCR*, illustrated using genomic data from the Ag1000G Phase 2 release. To estimate the probability that a doubleton composed of two cytosines (C's), occurring at a site where the major allele is an adenosine (A), is recurrent, we first estimate the number of doubletons of this type that we expect to be the result of multiple mutations at the same site. There are 10,620 $A \to C, T$; 8,640 $A \to C, G$; and 22,938 $A \to T, G$ triallelic singletons (Table 1, Main Text). Following Eq. (2) in the main text, we multiply the number of observed triallelic singletons where one of the alternative alleles is a result of our mutation of interest ($A \to C$), and divide by twice the number of triallelic sites containing our major allele, A, but not our nucleotide of interest, C. Thus we arrive at:

$$N_{est}[A \to C, C] = \frac{10,620 \times 8,640}{2 \times 22,938}$$

$$= 2,002$$

This gives the estimated number of $A \to C$ doubletons that are the result of two independent $A \to C$ mutations. As there are a total of 40,394 $A \to C$ doubletons, we therefore estimate that $2,002/40,394 = 5\%$ of them are the result of recurrent mutation (Table 2, Main Text).

### Derivation of Eq. (5) in Main text

We wish to derive an expression for the conditional probability that a doubleton ($d$) with a linked rare variant ($LRV$) identifies sequences that are reciprocal closest relatives ($RCR$) at that site, $P[RCR|d, LRV]$. Using Bayes' theorem for three events, and showing each step:

$$\begin{aligned} P[RCR|d, LRV] &= \frac{P[RCR, d, LRV]}{P[d, LRV]} \\ &= \frac{P[LRV|d, RCR]\ P[d, RCR]}{P[d, LRV]} \\ &= \frac{P[LRV|d, RCR]\ P[RCR|d]\ P[d]}{P[LRV|d]\ P[d]} \\ &= \frac{P[LRV|d, RCR]\ P[RCR|d]}{P[LRV|d]} \end{aligned} \tag{SI-4}$$

$P[LRV|d]$, the proportion of doubletons having an $LRV$, is gotten by summing the two relevant pathways in the tree diagram (Fig. SI-1), combining those that are $RCR$ and those that are not:

$$P[LRV|d] = P[LRV|d, RCR]\ P[RCR|d] + P[LRV|d, \neg RCR]\ (1 - P[RCR|d]) \tag{SI-5}$$

Similarly, the proportion of non-doubletons having an $LRV$ is:

$$P[LRV|\neg d] = P[LRV|\neg d, RCR]\ P[RCR|\neg d] + P[LRV|\neg d, \neg RCR]\ (1 - P[RCR|\neg d]) \tag{SI-6}$$

We next define $k_1$ as the ratio of the probabilities that $RCR$ sequences with and without a doubleton

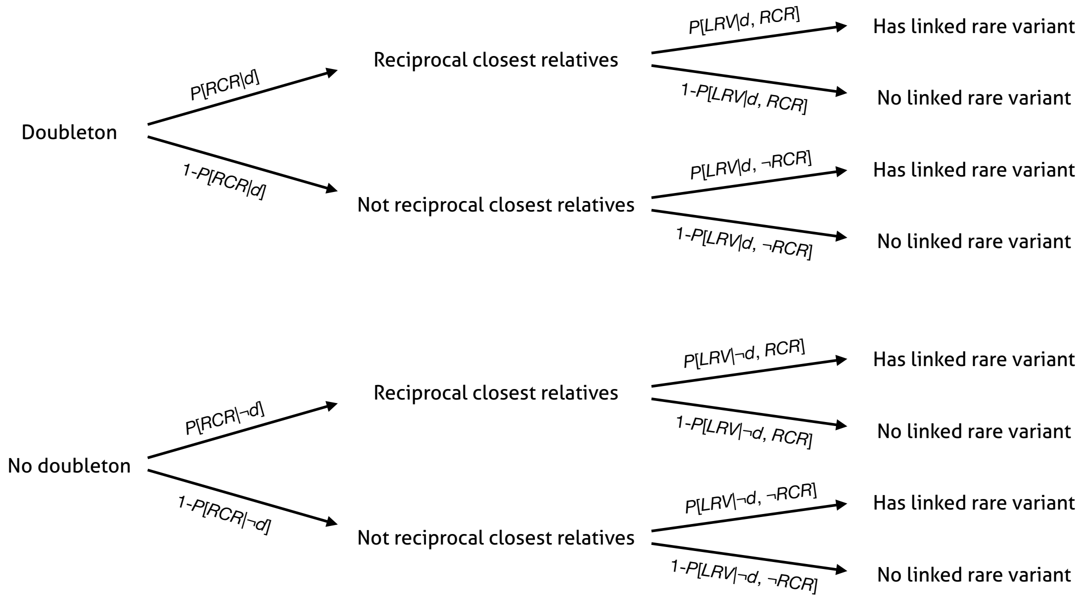


Figure SI-1: Diagrammatic representation of the probability comparisons used. Comparisons are made between doubleton and random pairs; for each set, we distinguish between those which are reciprocal closest relatives ($RCR$) and non-$RCR$, and then whether these carry a given linked rare variant ($LRV$) or not. The combined probabilities of these (tips) can be calculated from empirical observations of population genomic data, and combined to deduce unknown values.

have an $LRV$, and $k_2$ as the same ratio for non-$RCR$ sequences and make the substitutions

$$P[LRV|d, RCR] = k_1 P[LRV|\neg d, RCR] \qquad \text{and} \qquad P[LRV|d, \neg RCR] = k_2 P[LRV|\neg d, \neg RCR] \tag{SI-7}$$

into Equation (SI-5). We then simplify the notation and re-write Equations (SI-5 & SI-6) as:

$$a = k_1 bx + k_2(1-b)y \qquad \text{and} \qquad c = dx + (1-d)y \tag{SI-8}$$

where

$$\begin{aligned} a &= P[LRV|d] \\ b &= P[RCR|d] \\ c &= P[LRV|\neg d] \\ d &= P[RCR|\neg d] \\ x &= P[LRV|\neg d, RCR] \\ y &= P[LRV|\neg d, \neg RCR] \end{aligned}$$

Equations (SI-8) can then be solved to give

$$x = \frac{a(1-d) - k_2(1-b)c}{k_1 b(1-d) - k_2 d(1-b)} \tag{SI-9}$$

$$y = \frac{k_1 bc - ad}{k_1 b(1-d) - k_2 d(1-b)} \tag{SI-10}$$

Substituting Equation (SI-9) into (SI-7) and then (SI-4), we get:

$$P[RCR|d, LRV] = \frac{k_1\ b\ (a\ (1-d) - k_2\ c\ (1-b))}{a\ (k_1\ b\ (1-d) - k_2\ d\ (1-b))} \tag{SI-11}$$

The four conditional probabilities on the right can be estimated as detailed in the main text, leaving only $k_1$ and $k_2$ to be considered. $k_1$ is the answer to the question, "How much more likely are two sequences that are $RCR$ at a site to have an $LRV$ if the site is a doubleton than if it is not." This is a variant of the "inspection paradox" (see also Mathieson and McVean (2014)). If two sequences are $RCR$, the probability they have a doubleton will depend on the length of the branch leading from their common ancestor back to the previous coalescent event — longer branches are more likely to have had a mutation on them than shorter branches, and therefore $RCR$ doubletons will on average be associated with longer branches leading to that common ancestor than $RCR$ non-doubletons. Linked sites will tend to have a similar genealogical history, and therefore will also have an increased probability of having a doubleton. Therefore, we expect $k_1 > 1$. If we assume those branch lengths are approximately exponentially distributed, then branches that have a mutation on them will be, on average, twice the length of those that do not, and therefore, if we restricted $LRV$s to be doubletons, we would expect $k_1 \approx 2$. However, the effect will be smaller for $LRV$s that are tripletons, quadrupletons, etc., the majority of which are expected to be due to mutations on another, higher branch on the genealogy. Therefore, overall, we expect $1 < k_1 < 2$. A similar logic should apply to $k_2$, which quantifies how much more likely are two sequences that are not $RCR$ at a site to share an $LRV$ if the site is a doubleton than if it is not. Non-$RCR$ doubletons are due to recurrent singleton mutations, and are more likely to occur on long external branches than on short ones; linked sites will have a similar genealogical history, and be more likely to also have recurrent mutations, but the effect will be weaker for tripletons, quadrupletons, etc., so overall we again expect $1 < k_2 < 2$. Setting either $k_1$ or $k_2$ equal to 2 instead of 1 gives a slightly lower probability $P[RCR|d, LRV]$, and therefore is more conservative; in practise, with our data the difference is minimal: for our threshold definition of an $LRV$ to be one that is found in 2-18 sequences in our dataset, $P[RCR|d, LRV] = 99.3\%$ for $k_1 = k_2 = 1$ and $98.6\%$ for $k_1 = k_2 = 2$. In principle, complex mutational events that lead to simultaneous changes at more than one site could also affect $k_1$ and $k_2$ if $LRV$s were restricted to doubletons, but again any such effect should be considerably weaker for higher frequency $LRV$s.

Setting $k_1 = k_2 = 1$ in Equation (SI-11) gives:

$$P[RCR|d, LRV] = \frac{b\ (a\ (1-d) -\ c\ (1-b))}{a\ (b-d)} \tag{SI-12}$$

which was used to produce Figure 1B in the main text. The Bayes Factor is then calculated as the ratio of the posterior odds to the prior odds:

$$BF = \frac{PosteriorOdds}{PriorOdds}$$

$$= \frac{\frac{P[RCR|d,LRV]}{1-P[RCR|d,LRV]}}{\frac{P[RCR|d]}{1-P[RCR|d]}}$$

$$= 1 + \frac{a-c}{bc-ad}$$

or, using the more explicit notation:

$$BF = 1 + \frac{P[LRV|d] - P[LRV|\neg d]}{P[RCR|d]\ P[LRV|\neg d] - P[LRV|d]\ P[RCR|\neg d]} \tag{SI-13}$$

as given in the main text.

As an aside, we note that a similar approach can be taken to derive an expression for the probability a doubleton has separate origins and the sequences are not $RCR$ when there is no $LRV$ as a function of the same four observables:

$$P[\neg RCR|d,\neg LRV] = \frac{P[\neg RCR,d,\neg LRV]}{P[d,\neg LRV]}$$

$$= \frac{P[\neg LRV|d,\neg RCR]\ P[d,\neg RCR]}{P[d,\neg LRV]}$$

$$= \frac{P[\neg LRV|d,\neg RCR]\ P[\neg RCR|d]\ P[d]}{P[\neg LRV|d]\ P[d]}$$

$$= \frac{P[\neg LRV|d,\neg RCR]\ P[\neg RCR|d]}{P[\neg LRV|d]}$$

$$= \frac{(1-P[LRV|d,\neg RCR])\ (1-P[RCR|d])}{1-P[LRV|d]} \tag{SI-14}$$

Using Equations (SI-10 & SI-7) this becomes:

$$P[\neg RCR|d, \neg LRV] = \frac{1 - P[RCR|d]}{1 - P[LRV|d]} \left(1 + \frac{k_2 P[LRV|d]\ P[RCR|\neg d] - k_1 k_2 P[RCR|d]\ P[LRV|\neg d]}{k_1 P[RCR|d]\ (1 - P[RCR|\neg d]) - k_2 P[RCR|\neg d]\ (1 - P[RCR|d])}\right) \tag{SI-15}$$

However, for our data these yield relatively low probabilities and so appear of little use for identifying a subset of doubletons with a high likelihood of having separate origins.
